# Discovery of highly potent small molecule pan-coronavirus fusion inhibitors

**DOI:** 10.1101/2023.01.17.524492

**Authors:** Francesca Curreli, Kent Chau, Thanh-Thuy Tran, Isabella Nicolau, Shahad Ahmed, Pujita Das, Christopher D. Hillyer, Mary Premenko-Lanier, Asim K. Debnath

## Abstract

The unprecedented pandemic of COVID-19, caused by a novel coronavirus, SARS-CoV-2, has led to massive human suffering, death, and economic devastation worldwide. The virus is mutating fast to more transmissible and infectious variants. The Delta variant (B.1.617.2), initially identified in India, and the omicron variant (BA.4 and BA.5) have spread worldwide. In addition, recently alarming antibody evasive SARS-CoV-2 subvariants, BQ and XBB, have been reported. These new variants may pose a substantial challenge to controlling the spread of this virus. Therefore, the continued development of novel drugs having pan-coronavirus inhibition to treat and prevent infection of COVID-19 is urgently needed. These drugs will be critically important in dealing with new pandemics that will emerge in the future. We report the discovery of several highly potent small molecule pan-coronavirus inhibitors. One of which, NBCoV63, showed low nM potency against SARS-CoV-2 (IC_50_: 55 nM), SARS-CoV (IC_50_: 59 nM), and MERS-CoV (IC_50_: 75 nM) in pseudovirus-based assays with excellent selectivity indices (SI: as high as > 900) demonstrating its pan-coronavirus inhibition. NBCoV63 showed equally effective antiviral potency against SARS-CoV-2 mutant (D614G) and several variants of concerns (VOCs) such as B.1.617.2 (Delta), B.1.1.529/BA.1 and BA.4/BA.5 (Omicron) and K417T/E484K/N501Y (Gamma). NBCoV63 also showed similar efficacy profiles to Remdesivir against authentic SARS-CoV-2 (Hong Kong strain) and two of its variants (Delta and Omicron) by plaque reduction in Calu3 cells. Additionally, we show that NBCoV63 inhibits virus-mediated cell-to-cell fusion in a dose-dependent manner. Furthermore, the Absorption, distribution, metabolism, and excretion (ADME) data of NBCoV63 demonstrated drug-like properties.

## INTRODUCTION

The devastating effect of coronavirus disease 2019 (COVID-19) caused by the novel SARS-CoV-2, first reported in December 2019 in Wuhan^1^, China, is continuing worldwide. Although two other coronaviruses (CoVs), SARS-CoV-1 in 2003 and MERS-CoV in 2012, were responsible for two severe outbreaks, the current COVID-19 pandemic outpaced all other known pandemics to date. According to data from the Johns Hopkins Coronavirus Resource Center, as of January 17, 2022, there were more than 667 million cases of COVID-19 and 6.7 million deaths globally. In the US, over 101 million cases and more than 1.1 million deaths have been reported (https://coronavirus.jhu.edu/map.html).

Although the massive death, hospitalization, uncertainty, and panic have subsided considerably, the continuous emergence of mutant variants with more powerful transmissibility and evading capacity to the currently available vaccines is causing great concern. Furthermore, there have been many reports of “breakthrough” SARS-CoV-2 infections among people who were fully vaccinated^2, 3^. In addition, a recent report indicates that about 25% of the world population is hesitant to vaccinate against SARS-CoV-2^4^. Currently, two small molecule drugs are approved by the FDA, Veklury (Remdesivir; Gilead, USA) and Olumiant (Baricitinib, Lilly, USA). In addition, two small molecule drugs, Paxlovid (a combination of Nirmatrelvir and Ritonavir; Pfizer, USA) and Lagevrio (Molnupiravir; Merck, USA), are approved under an emergency use authorization (EUA). Unfortunately, the resistant mutants of Remdesivir^5, 6^, and Paxlovid^7^ have recently been reported. Furthermore, most approved vaccines and antibody-based therapies show substantial loss of potency against SARS-CoV-2 variants of concern (VOCs)^8-11^. The Delta variant (B.1.617.2)^3, 12, 13^, initially identified in India, and the Omicron variant (BA.4 and BA.5) have spread worldwide. Recently, David Ho and his team reported alarming antibody evasive SARS-CoV-2 subvariants, BQ and XBB^14^. These new variants may pose a substantial challenge to controlling the spread of this virus. Therefore, the continued development of novel drugs having pan-coronavirus inhibition to treat and prevent infection of COVID-19 is urgently needed.

Coronaviruses are positive-sense, enveloped, single-stranded RNA viruses in the family Coronaviridae. The life cycle of these enveloped viruses begins by attaching the trimeric spike surface protein (S) to a host cell ^15, 16^, which is then cleaved into S1 and S2 subunits by furin-like proteases. The S1 subunit of the S protein uses its receptor-binding domain (RBD) to bind to a host-cell receptor, known as the angiotensin-converting enzyme 2 (ACE2)^17, 18^. SARS-CoV and SAR-CoV-2 use the same cellular receptor and the cellular transmembrane serine protease 2 (TMPRSS2) for the S protein priming^19-21^. The receptor for MERS-CoV is dipeptidyl peptidase 4 (DPP4; CD26)^22^. The binding of the S1 subunit to a receptor triggers the fusion protein (FP) of the S2 subunit to insert into the cell membrane and help the heptad repeat region 1 (HR1) domain of S2 to form a coiled-coil trimer. This process initiates the binding of the heptad repeat region 2 (HR2) domain of S2 to a hydrophobic groove in the HR1 trimer in an antiparallel manner to form a six-helix bundle (6-HB) structure. The 6-HB formation is typical of other class I membrane fusion proteins in viruses such as influenza, human immunodeficiency virus (HIV), and Ebola^15, 23^. This brings the viral and host-cell membranes together for virus–cell fusion^24^, a critical step for virus entry into host cells. The fusion mechanism for the S protein of other CoVs, including MERS-CoV, is similar.

The S protein of CoVs is a surface protein and is exposed on the surface of the mature virus particle. Consequently, it is the primary target for developing neutralizing antibodies and vaccines. The S1 subunit, primarily the RBD domain, and the S2 subunit, especially the HR1 domain, have been targeted for novel drug design^25-33^. However, the RBD of the S1 domain is not well conserved among CoVs^34^. As a result, neutralizing antibodies of SARS-CoV show poor cross-reactivity with SARS-CoV-2^35, 36^.

Furthermore, the RBD domain of SARS-CoV-2 has been reported to exhibit a higher number of mutations^37-39^. Some of these mutations reduce antibodies’ efficacy, hence the currently available vaccines^40, 41^. Recently, BQ.1, BQ.1.1, XBB, and XBB.1 variants of SARS-CoV-2 with multiple mutations on the RBD have been reported, which markedly reduce serum neutralization^14^. Therefore, the RBD may not be an ideal target for developing novel pan-coronavirus inhibitors.

Contrary to the RBD domain, the membrane fusion domains located in the S2 subunit are primarily conserved, consequently, can be considered as a potential drug target for developing novel small molecule and peptide-based pan-coronavirus inhibitors. These drugs will be critically important in dealing with new pandemics that are certain to emerge in the future.

In this study, we report the discovery of a series of small molecule compounds with highly potent pan-coronavirus activity against SARS-CoV-2, SARS-CoV-1, and MERS-CoV.

## Results and Discussion

### Identification of pan-coronavirus inhibitors

An in-depth analysis of the postfusion hairpin structures of CoVs indicated that they were not only structurally similar but also shared critical salt bridges between the HR1 and HR2 regions^30^. For example, K947 of the HR1 domain in SARS-CoV-2 forms a salt bridge with E1182 of the HR2 domain. Similarly, K929 of the HR1 domain in SARS-CoV forms a salt bridge with E1163 of the HR2 domain. Furthermore, a salt bridge between K1021 with E1265 is also present in the MERS-CoV postfusion spike structure^42^. Interestingly, we reported a similar salt bridge interaction between K547 of the N-terminal heptad repeat region (also known as HR1) and D632 of the C-terminal heptad repeat region (also known as HR2) in the HIV-1 gp41 hairpin structure^43^. The later observation motivated us to design a series of highly potent benzoic acid-based inhibitors of HIV-1 gp41 fusion, which contain a COOH group^44^. Based on these remarkable similarities in the formation of salt bridges to form the 6-HB structures and their involvement in virus–cell fusion of HIV-1 gp41 and the CoV S proteins, we hypothesized that COOH-containing inhibitors take part in salt bridge formations by fitting into a cavity in the prefusion trimer structures of CoVs to prevent the 6-HB formation and subsequent virus–cell fusion.

Peptide-based pan-coronavirus fusion inhibitors were recently shown to possess potent inhibitory activity against HIV-1, HIV-2, and SIV^45^. These encouraging results motivated us to screen a set of COOH-containing molecules, which we referred to as NBCoVs and tested them against SARS-CoV-2, SARS-CoV, and MERS-CoV in spike-pseudotyped antiviral assays^46^. However, the NBCoV molecules contain an ene-rhodanine scaffold, a well-known frequent hitter, which suggests that they might be pan assay interference compounds (PAINS)^47-49^. Since the first publications on PAINS^50^ and colloidal aggregators^51^ as frequent hitters or promiscuous inhibitors in high-throughput screening, PAINS-containing compounds are generally considered “undevelopable”. However, many suggest that all frequent hitters should not be discarded without validating whether they are target-specific or truly promiscuous^52-56^. One of the strong counterarguments was put forward by Bajorath, who argued that the chemical integrity and specific biological activity of compounds containing PAINS scaffolds should be considered in the context of the overall larger structure^53^. These controversies and counterarguments led nine American Chemical Society (ACS) editors to recommend the necessary steps to rule out any artifacts from PAINS-containing compounds in biological activities^57^. Even though we followed the recommendations of the ACS editors^57^ and confirmed through concurrent and orthogonal experiments that NBCoV molecules are not promiscuous and aggregators^46^, we decided not to pursue those leads to avoid any future controversy. Instead, we were confident that despite the presence of a PAINS scaffold in these molecules, their antiviral activity must be due to the distinct pharmacophores of the molecules. We conclusively established the structure-activity relationship (SAR). We demonstrated that despite the presence of the same ene-rhodanine moiety in the compounds used as controls, they showed no antiviral activity at all^46^. We were confident about the SAR information from that study and used that to search compounds from commercial small molecule drug-like databases. We identified more than 20 molecules that did not exhibit any PAINS^50, 56, 58^ alerts when run through the SwissADME^59^ server.

### Antiviral activity and cytotoxicity of the NBCoV small molecules in a pseudovirus assay

The anti-coronavirus activity of the newly identified NBCoV small molecules was evaluated by infecting two cell types: 293T/ACE2 cells, overexpressing the human receptor ACE2 and A549/ACE2/TMPRSS2 (A549/AT) cells overexpressing both, ACE2 and TMPRSS2. Cells were infected with aliquots of the SARS-CoV-2 (USA-WA1/2020) pseudovirus pretreated with escalating concentrations of the NBCoV small molecules for 30 minutes to calculate the concentration required to inhibit 50% (IC_50_) of SARS-CoV-2 infection. In parallel, the cytotoxicity of the compounds was evaluated in both cell lines to calculate the CC_50_ (the concentration for 50% cytotoxicity), and the Selectivity Indexes (SI) (Table 1). As shown, compound NBCoV63 had an IC_50_ of 80±7 and 55±3 nM, respectively, and CC_50_ ≥50 μM (SI of >625 and 909 for the two cell lines), which was the highest dose we could test for all the compounds due to poor solubility. NBCoV35 and NBCoV37, which have a similar structure to NBCoV63, also showed low nanomolar activity in both cell lines (IC_50_ = 66-198 nM). While NBCoV35 had similar toxicity in both cell lines, NBCoV37 induced higher toxicity in 293T/ACE2 cells. NBCoV36 antiviral activity was slightly weaker than the previous three compounds (330±18 and 227±18 nM); we noticed in this case as well that NBCoV36 exhibited higher toxicity in 293T/ACE2 cells than in A549/AT. NBCoV81 also had an antiviral activity with SI >109 in both cell lines. The remaining compounds had minor to no activity. Based on the results reported in **Table 1**, we decided to validate further the anti-coronavirus activity of the compounds that showed the best SI (NBCoV35-37 and NBCoV63) against SARS-CoV2 VOCs and other coronaviruses in neutralization assays.

**Table 1.**
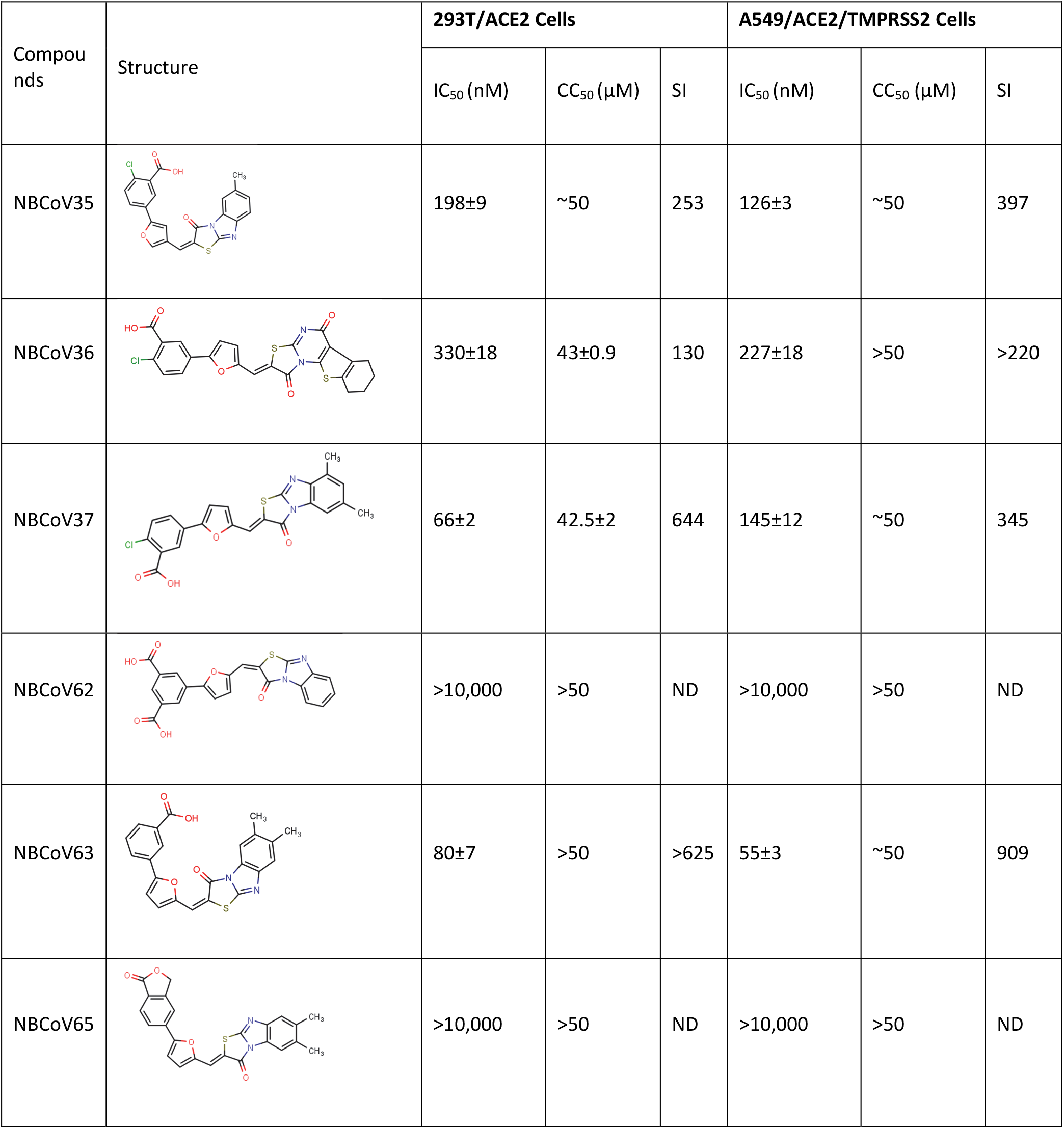
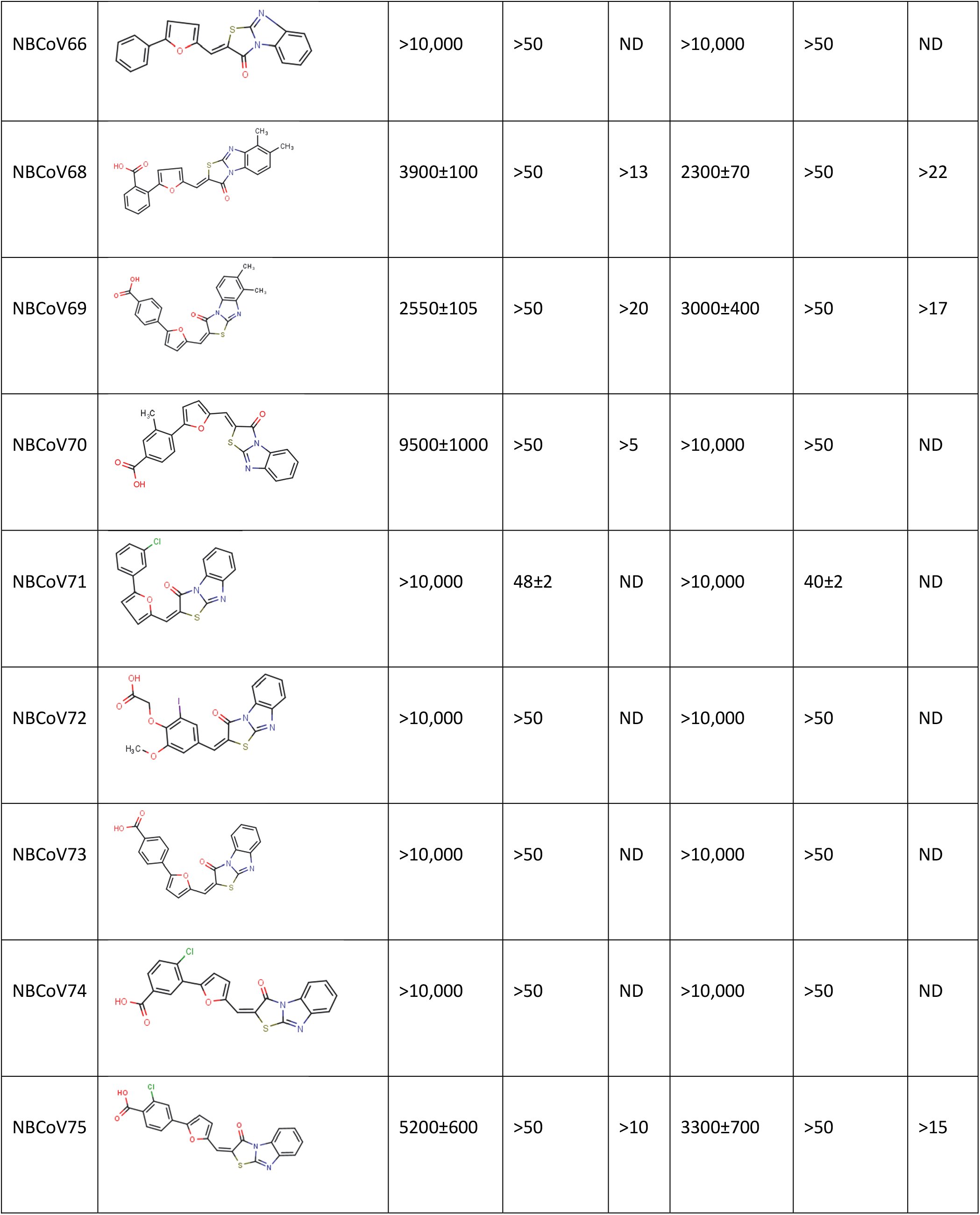
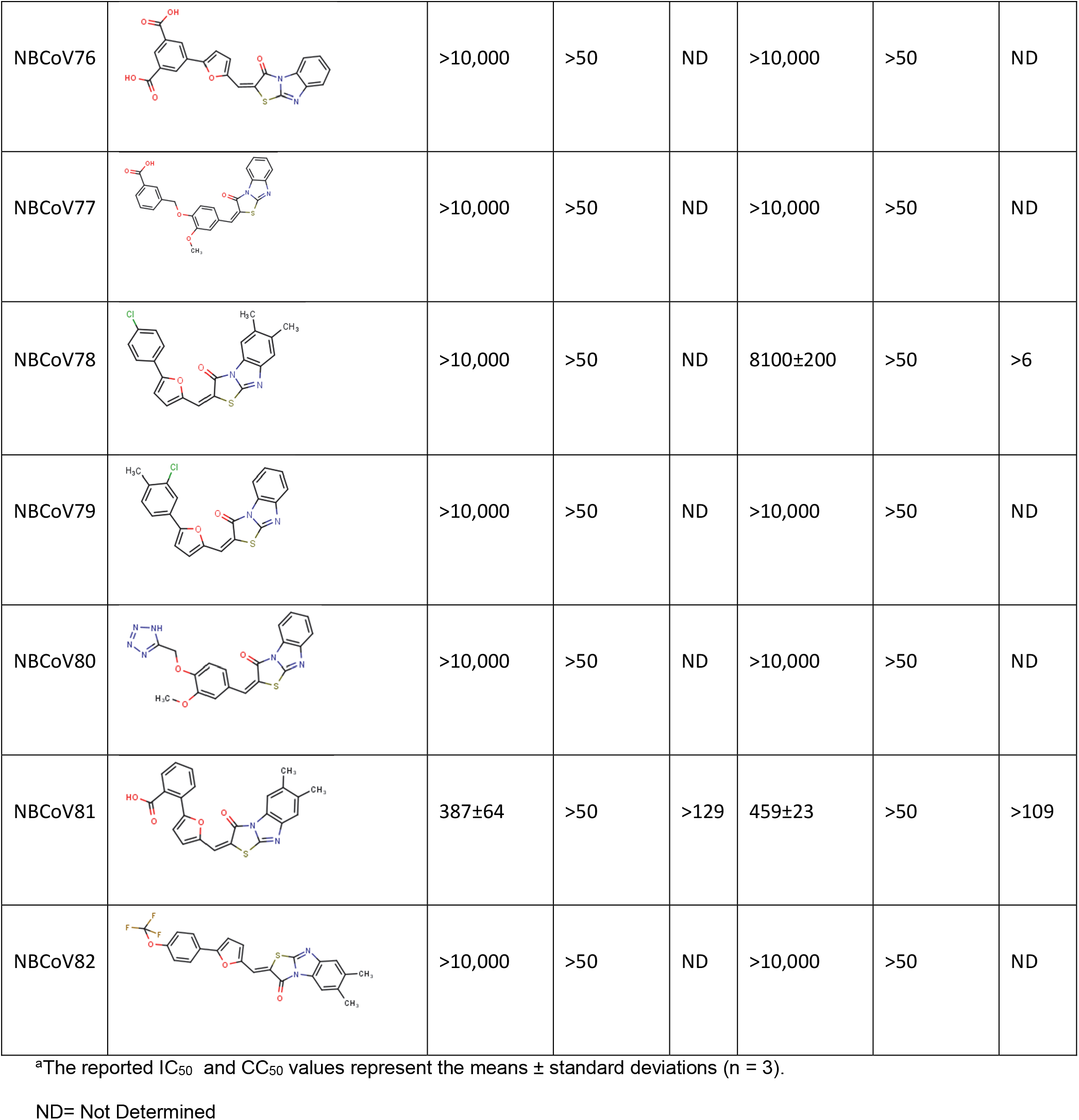
Screening of the neutralization activity of NBCoV compounds against NL4-3^ΔEnv^-NanoLuc/SARS-CoV-2 (USA-WA1/2020) pseudovirus (IC_50_), toxicity evaluation (CC_50_) and Selectivity Indexes (SI) in two cell lines.

| Compounds | Structure | 293T/ACE2 Cells |  |  | A549/ACE2/TMPRSS2 Cells |  |  |
| --- | --- | --- | --- | --- | --- | --- | --- |
|  |  | IC<sub>50</sub> (nM) | CC<sub>50</sub> (μM) | SI | IC<sub>50</sub> (nM) | CC<sub>50</sub> (μM) | SI |
| NBCoV35 |  | 198±9 | ~50 | 253 | 126±3 | ~50 | 397 |
| NBCoV36 |  | 330±18 | 43±0.9 | 130 | 227±18 | >50 | >220 |
| NBCoV37 |  | 66±2 | 42.5±2 | 644 | 145±12 | ~50 | 345 |
| NBCoV62 |  | >10,000 | >50 | ND | >10,000 | >50 | ND |
| NBCoV63 |  | 80±7 | >50 | >625 | 55±3 | ~50 | 909 |
| NBCoV65 |  | >10,000 | >50 | ND | >10,000 | >50 | ND |
| NBCoV66 |  | >10,000 | >50 | ND | >10,000 | >50 | ND |
| NBCoV68 |  | 3900±100 | >50 | >13 | 2300±70 | >50 | >22 |
| NBCoV69 |  | 2550±105 | >50 | >20 | 3000±400 | >50 | >17 |
| NBCoV70 |  | 9500±1000 | >50 | >5 | >10,000 | >50 | ND |
| NBCoV71 |  | >10,000 | 48±2 | ND | >10,000 | 40±2 | ND |
| NBCoV72 |  | >10,000 | >50 | ND | >10,000 | >50 | ND |
| NBCoV73 |  | >10,000 | >50 | ND | >10,000 | >50 | ND |
| NBCoV74 |  | >10,000 | >50 | ND | >10,000 | >50 | ND |
| NBCoV75 |  | 5200±600 | >50 | >10 | 3300±700 | >50 | >15 |
| NBCoV76 |  | >10,000 | >50 | ND | >10,000 | >50 | ND |
| NBCoV77 |  | >10,000 | >50 | ND | >10,000 | >50 | ND |
| NBCoV78 |  | >10,000 | >50 | ND | 8100±200 | >50 | >6 |
| NBCoV79 |  | >10,000 | >50 | ND | >10,000 | >50 | ND |
| NBCoV80 |  | >10,000 | >50 | ND | >10,000 | >50 | ND |
| NBCoV81 |  | 387±64 | >50 | >129 | 459±23 | >50 | >109 |
| NBCoV82 |  | >10,000 | >50 | ND | >10,000 | >50 | ND |

<sup>a</sup>The reported IC<sub>50</sub> and CC<sub>50</sub> values represent the means ± standard deviations (n = 3).
ND= Not Determined

### Antiviral activity of NBCoV35-37 and NBCoV63 against SARS-CoV-2 mutant (D614G) and four VOCs

SARS-CoV-2 VOCs carrying multiple mutations in their spike are the cause of concerns due to increased virulence and reduced vaccine efficacy. In this study, we decided to evaluate the effectiveness of our best NBCoV compounds against the first detected virus with a single spike mutation, USA-WA1/2020/D614G, and four VOCs already spread worldwide: Delta B.1.617.2, Omicron B.1.1.529/BA.1, Omicron BA.4/BA.5 and Brazil (Gamma, carrying three spike mutations: K417T/E484K/N501Y) (**Table 2**). We infected 293T/ACE2 cells and A549/AT cells with the mutant SARS-CoV-2 pseudoviruses in the absence or presence of the NBCoV compounds. We observed that NBCoV63 had potent antiviral activity against all variant pseudoviruses assessed, as indicated by the low IC_50_s detected, ranging from 34-96 nM in 293T/ACE2 cells and 26-105 nM in A549/AT cells (**Table 2**). The best activity of NBCoV35 was recorded against Omicron BA.4/BA.5 in both cell lines (IC_50_s: 94±6 and 170±13 nM, respectively), while it seemed weaker against the Brazil variant (IC_50_s: 506±44 and 292±23 nM). NBCoV37 was also active against all the variants, with IC_50_ values in the 82-267 nM range in 293T/ACE2 cells and 42.5-311 nM in A549/AT cells. Finally, NBCoV36 displayed lower antiviral activity exhibiting a wider range of IC_50_ values (181-1183 nM in 293T/ACE2 cells and 129-332 nM in A549/AT cells). These results suggest that the NBCoV small molecules maintain their antiviral activity against the SARS-CoV-2 D614G-mutant and other VOCs.

**Table 2.** Antiviral activity of the NBCoV small molecules in the single-cycle assay in two cell lines against pseudovirus NL4-3^ΔEnv^-NanoLuc/SARS-CoV-2 mutant (D614G) and its variants of concern (VOC) (IC_50_).

| Compounds | SARS-CoV-2 |  |  |  |  |
| --- | --- | --- | --- | --- | --- |
|  | IC<sub>50</sub> (nM)<sup>a</sup> |  |  |  |  |
|  | USA-WA1/2020<br>D614G | Variants of Concern (VOC) |  |  |  |
|  |  | Delta<br>(B.1.617.2) | Omicron<br>(B.1.1.529/BA.1) | Omicron<br>(BA.4/BA.5) | Brazil (Gamma:<br>K417T/E484K/N501Y) |
| 293T/ACE2 Cells |  |  |  |  |  |
| NBCoV35 | 195±10 | 335±5 | 173±30 | 94±6 | 506±44 |
| NBCoV36 | 430±31 | 802±143 | 368±90 | 181±12 | 1183±58 |
| NBCoV37 | 157±1.5 | 267±23 | 135±3 | 82±8 | 260±52 |
| NBCoV63 | 40.8±1.2 | 84±0.5 | 96±11 | 34±1 | 77.5±17.3 |
| **A549/ACE2/TMPRSS2 Cells** |  |  |  |  |  |
| NBCoV35 | 266±15 | 236±6 | 201±38 | 170±13 | 292±23 |
| NBCoV36 | 150±2 | 332±26 | 129±2 | 310±21 | 149±26 |
| NBCoV37 | 311±33 | 130±35 | 42.5±1 | 215±10 | 73±14 |
| NBCoV63 | 42.5±1 | 105±1 | 60±3 | 26±0.5 | 45.5±2 |

<sup>a</sup>The reported IC<sub>50</sub> values represent the means ± standard deviations (n = 3).

To further validate our findings, we evaluated the compounds in another human lung cell line, Calu-3. These non-small-cell lung cancer cells are permissive to SARS-CoV-2 strains, SARS-COV, and MERS virus strains. These cells were infected with SARS-CoV2 (USA-WA1/2020) and the most recent VOC, Omicron BA.4/BA.5. Data reported in **Table 3** shows that NBCoV63 maintained its excellent antiviral activity against both viral clones assessed with IC_50_ of about 120 nM. On the contrary, NBCoV35-37 showed higher IC_50_s in Calu-3 cells than those detected with the other cell lines. However, they maintained good antiviral activity against the omicron BA.4/BA.5 variants. The CC_50_ in these cells suggested no interference from toxicity.

**Table 3.** Antiviral activity of the NBCoV small molecules in the single-cycle assay in Calu-3 cells against pseudovirus NL4-3^ΔEnv^-NanoLuc/SARS-CoV-2 (USA-WA1/2020) and the VOC Omicron (BA.4/BA.5) (IC_50_), toxicity (CC_50_) and SI.

| Compounds | Calu-3 Cells |  |  |  |  |
| --- | --- | --- | --- | --- | --- |
|  | SARS-CoV-2 |  |  |  |  |
|  | USA-WA1/2020 |  | Omicron BA.4/BA.5 |  | CC<sub>50</sub> |
|  | IC<sub>50</sub> (nM)<sup>a</sup> | SI | IC<sub>50</sub> (nM)<sup>a</sup> | SI | (μM)<sup>a</sup> |
| NBCoV35 | 360±35 | >139 | 520±27 | >96 | >50 |
| NBCoV36 | 433±49 | >116 | 565±79 | >89 | >50 |
| NBCoV37 | 368±45 | >136 | 583±35 | >86 | >50 |
| NBCoV63 | 120±6 | 417 | 126±5 | 397 | 50 |

<sup>a</sup>The reported IC<sub>50</sub> and CC<sub>50</sub> represent the means ± standard deviations (n = 3).

To evaluate these small molecules as pancoronavirus inhibitors, we assessed them against the SARS-CoV (**Table 4**) and MERS-CoV (**Table 5**) pseudoviruses. We found that NBCoV63 was again the most potent compound in both cell lines infected with SARS-CoV (**Table 4**), with calculated IC_50_s of 110±0.5 nM (SI: 455) in 293T/ACE2 cells and 59±1.5 nM (SI: 848) in A549/AT cells. NBCoV35-37 also maintained their inhibitory activity against this virus with IC_50_ of 340-660 nM (SI: 65-132) in 293T/ACE2 cells and IC_50_ of 95-363 nM (SI: 138-526) in A549/AT cells.

**Table 4.** Antiviral activity of the NBCoV small molecules in a single-cycle assay in different cell lines against pseudovirus NL4-3^ΔEnv^-NanoLuc/SARS-CoV (IC_50_) and SI.

| Compound | SARS-CoV |  |  |  |
| --- | --- | --- | --- | --- |
|  | 293T/ACE2 Cells |  | A549/ACE2/TMPRSS2 Cells |  |
|  | IC<sub>50</sub> (nM)* | SI | IC<sub>50</sub> (nM)* | SI |
| NBCoV35 | 380±69 | 132 | 363±11 | 138 |
| NBCoV36 | 660±30 | 65 | 147±19 | 340 |
| NBCoV37 | 340±34 | 125 | 95±1 | 526 |
| NBCoV63 | 110±0.5 | 455 | 59±1.5 | 848 |

\*The reported IC<sub>50</sub> values represent the means ± standard deviations (n = 3).

**Table 5.** Antiviral activity of the NBCoV small molecules in the single-cycle assay in different cell lines against pseudovirus NL4-3^ΔEnv^-NanoLuc/MERS-CoV (IC^50^), toxicity (CC_50_) and SI.

| Compound | MERS-CoV |  |  |  |  |  |
| --- | --- | --- | --- | --- | --- | --- |
|  | Caco-2 Cells |  |  | MRC-5 Cells |  |  |
|  | IC<sub>50</sub> (nM)<sup>a</sup> | CC<sub>50</sub> (μM)<sup>a</sup> | SI | IC<sub>50</sub> (nM)<sup>a</sup> | CC<sub>50</sub> (μM)<sup>a</sup> | SI |
| NBCoV35 | 472±43 | >50 | >106 | 715±43 | >50 | >70 |
| NBCoV36 | 428±34 | >50 | >117 | 837±38 | >50 | >60 |
| NBCoV37 | 161±53 | >50 | >311 | 229±20 | 48±2 | 210 |
| NBCoV63 | 75±25 | >50 | >667 | 102±1 | >50 | >490 |

<sup>a</sup>The reported IC<sub>50</sub> and CC<sub>50</sub> values represent the means ± standard deviations (n = 3).

Furthermore, as reported in **Table 5**, we found that, yet again, NBCoV63 was the most potent compound against the MERS-CoV pseudovirus in both cell lines (IC_50_: 75 nM in Caco-2 cells, IC_50_: 102 nM in MRC-5 cells and SIs >667 and >490, respectively). In addition, NBCoV37 also had a potent MERS-CoV inhibitory activity with IC_50_ in the range 161-229 nM, while NBCoV35-36 had slightly weaker activity against this virus in both cell lines (IC_50_ 428-837). The data suggest that NBCoV35-37 and NBCoV63 indeed possess pancoronavirus activity.

Finally, to verify the specificity of the NBCoV compounds for the coronaviruses, we evaluated them against the amphotropic murine leukemia virus (A-MLV), which enters the cells via macropinocytosis^60^. We found that while NBCoV35-37 had insignificant activity against this virus, NBCoV63 showed instead some activity which was about 3.8-fold higher than the IC_50_ value detected against SARS-CoV-2 in the same cell line **(Table 6)**. Although further investigation will be necessary to explain this finding, we can exclude unspecific activity against the cells as the CC_50_ was >50 μM. Additionally, we performed two sets of experiments. In the first one, we exposed A549/AT cells to the compounds for 30 minutes before infection with SARS-CoV-2, and in the second experiment, we added the virus and the compounds to the cells simultaneously.

**Table 6.** Antiviral activity of the NBCoV compounds in a single-cycle assay in 293T/ACE2 cells against pseudovirus NL4-3^ΔEnv^Luc/A-MLV.

| Compound → | NBCoV35 | NBCoV36 | NBCoV37 | NBCoV63 |
| --- | --- | --- | --- | --- |
| Control Virus ↓ | IC<sub>50</sub> (nM)<sup>a</sup> |  |  |  |
| AMLV-NL4-3 | 1825±25 | 3300±100 | 1511±88 | 303±29 |

<sup>a</sup>The reported values represent the means ± standard deviations (n = 3).

We found no toxicity and no activity of the compounds against SARS-CoV-2 in both experiments (data not shown). These findings confirmed that the NBCoV compounds target the viruses to inhibit viral entry and do not affect the cell well-being as they did not induce cell toxicity and did not prevent cell infection when the cells were pretreated with the compounds, but they only inhibit viral infection when the virus was pretreated before adding it to the cells.

### NBCoV63 inhibited the replication-competent authentic (live) virus SARS-CoV-2 and two of its variants

Following the demonstration of the antiviral activity of NBCoV35 and NBCoV63 against mutant SARS-CoV-2 and VOCs, we next selected NBCoV63 as the best inhibitor as a lead compound. We tested its antiviral activity using a traditional plaque reduction assay against live SARS-CoV-2 variants. Three variants were selected SARS-CoV-2 Hong Kong/VM20001061/2020 tested at an 0.5 MOI, SARS-CoV-2 Omicron hCoV-19/USA/NY-MSHSPSP-PV56475/2022 and SARS-CoV-2 Delta hCoV-19/USA/PHC658/2021 both tested at 0.01 MOI. The control compound Remdesivir had an IC_50_ of 3.44 and 2.82 against the parent HK strain and the Omicron variant, respectively (**Table 7**), and an IC_50_ of 12.5 against the Delta variant. NBCoV63 had IC_50_ values that were lower for Delta and HK but slightly higher for the Omicron variant. There was no toxicity in Calu3 cells at the dilutions tested. These data demonstrate that NBCoV63 has similar antiviral activity as Remdesivir against live SARS-CoV-2 viruses.

**Table 7.** Antiviral activity against authentic viruses^1^

| Virus | Cells | IC<sub>50</sub> (μM)<sup>2</sup> |  | CC<sub>50</sub> (μM)<sup>3</sup> |
| --- | --- | --- | --- | --- |
|  |  | NBCoV63 | Remdesivir (control) |  |
| HK | Calu-3 | 2.04 | 3.44 | >50 |
| Delta (B.1.617.2) | Calu-3 | 9.27 | 12.5 |  |
| Omicron (BA.2.12.1) | Calu-3 | 5.45 | 2.82 |  |

<sup>1</sup> The assay was performed by SRI International, Menlo Park, CA
<sup>2</sup> The reported values represent the means ± standard deviations (n = 2) and the assay was repeated.
<sup>3</sup> CC<sub>50</sub> of both NBCoV63 and Remdesivir in Calu-3 cells was >50 μM.

### NBCoV small molecules inhibited the SARS-CoV-2 mediated cell-to-cell fusion

It has been reported that cell-to-cell direct contact/fusion is another efficient viral spreading mode^61^. It facilitates the infection of adjacent cells without producing free viruses and, at the same time, contributes to tissue damage and syncytia formation. We previously described a novel cell-to-cell fusion inhibition assay using Jurkat cells expressing a luciferase gene and the wild-type S gene from the SARS-CoV-2 Wuhan-Hu-1 isolate as donor cells and 293T/ACE2 cells as acceptor cells^46^. We used 293T/ACE2 cells cultured with or without Jurkat cells as a positive and negative control, respectively. As an additional control, we included 293T/ACE2 cells cultured with Jurkat cells expressing the luciferase gene only (Jurkat-Luc). We evaluated the inhibitory activity of escalating concentrations of NBCoV63 on cell-to-cell fusion (**Figure 1**). We found that NBCoV63 inhibits SARS-CoV-2–mediated cell-to-cell fusion in a dose-dependent manner, suggesting that this compound disrupts the binding of SARS-CoV-2 S protein with the receptor and interferes with virus-mediated cell-to-cell fusion.

**Figure 1.**
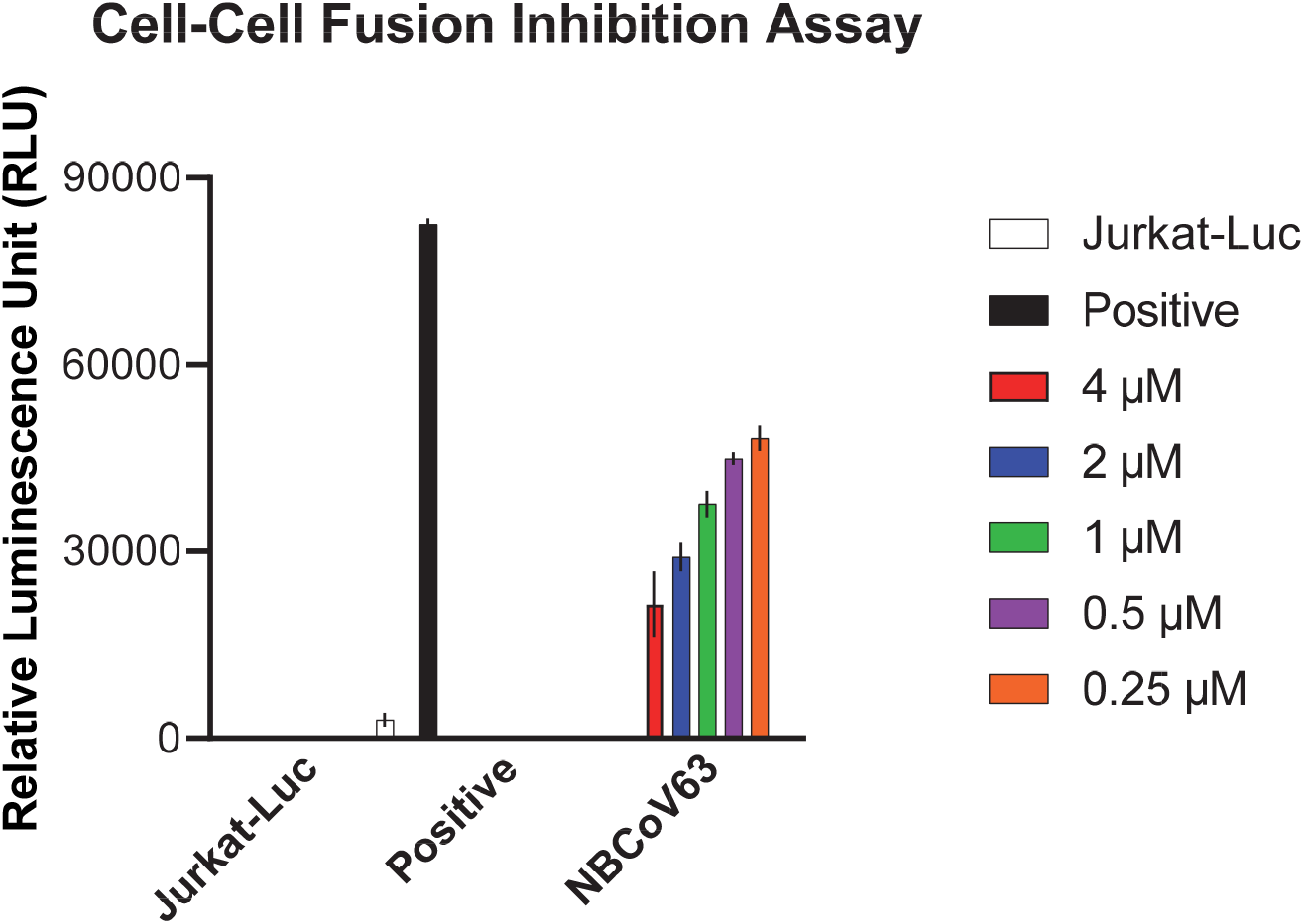
SARS-CoV-2-mediated cell-to-cell fusion inhibition by NBCoV63

### Docking study indicates NBCoV compounds bind to HR1 domain of SARS-CoV-2

Without any information about the structures of the inhibitors bound to the SARS-CoV-2 S protein, we performed a Glide-based docking study (Schrodinger, CA) using the SARS-CoV-2 prefusion form of the S protein (PDB: 6VSB) with an active-compound (NBCoV63) and an inactive-compound (NBCoV66) (**Figure 2**). We observed that the COOH moiety of NBCoV63 formed a salt bridge/H-bond with K947 of the A chain of the HR1 domain of the S protein (**Figure 2A**). Furthermore, we found that K776 of the B chain of the HR1 domain created similar salt bridge/H-bond interactions with the COOH of NBCoV63. **Figure 2B** confirms the importance of the COOH group at the meta position of the phenyl ring in the antiviral activity of this class of compounds. On the other hand, NBCoV66, which has all the critical scaffold in its structure but has no COOH group on the meta position of the phenyl group, showed no antiviral activity confirming the vital role the meta COOH group plays.

**Figure 2.**
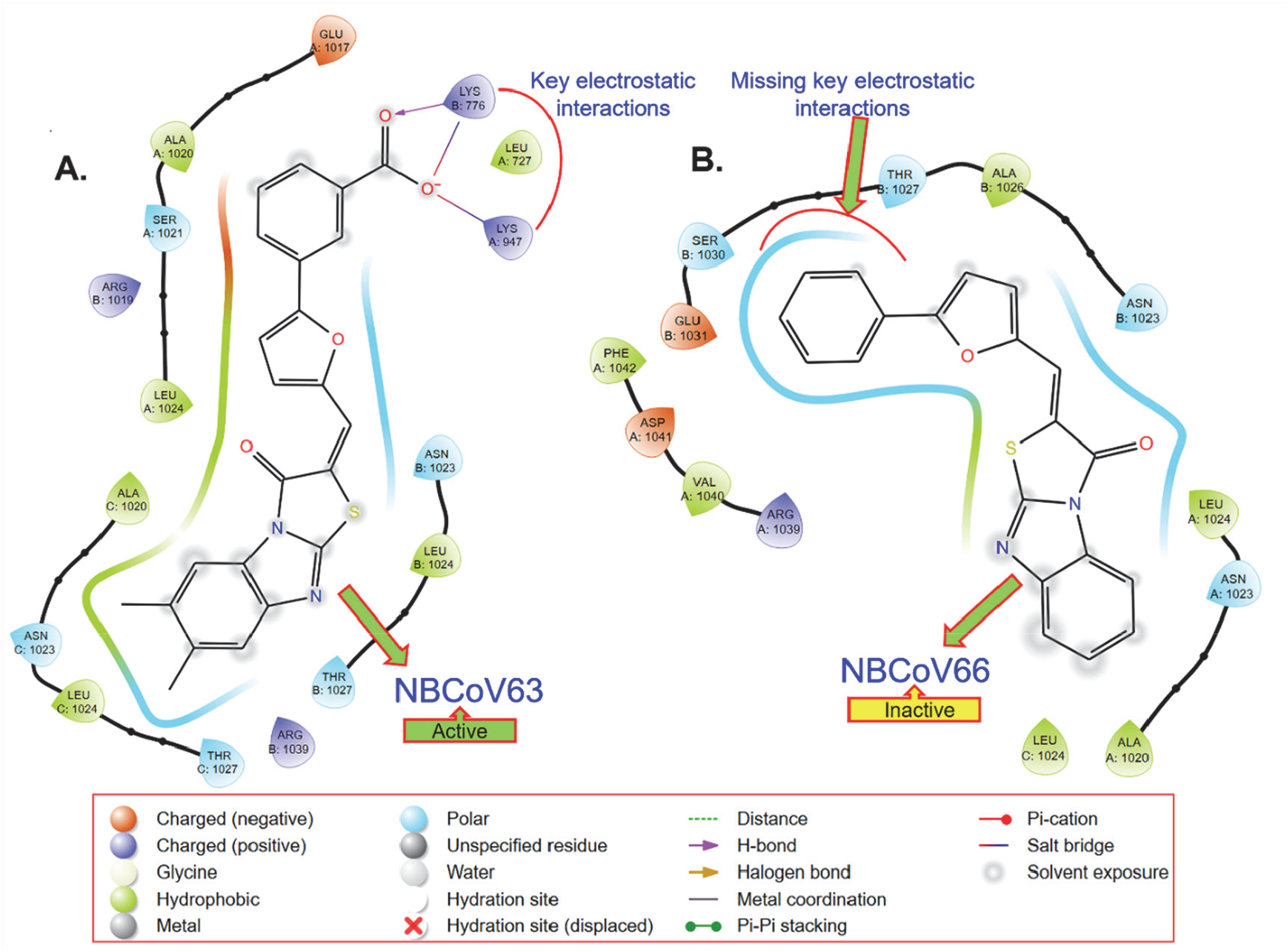
Glide-based docking of an active (NBCoV63) and an inactive (NBCoV66) demonstrate that (A) the COOH group of the active compound formed the key salt-bridge interaction; (B) this key interaction is missing in NBCoV66 due to the absence of the COOH group.

To validate the findings from the docking study, we reasoned that substituting K947 of the A chain and K776 of the B chain of the HR1 domain of the S protein would prevent the salt bridge/H-bond formation and, as a result, a loss of antiviral activity. To this end, we prepared a set of four mutant SARS-CoV-2 pseudoviruses carrying single mutations K947D, K947L, K776D, or K776L to assess the antiviral activity of the NBCoV molecules. We found that NBCoV36 and NBCoV37 completely lost their antiviral activity at the higher dose used (4000 nM) against all the mutant viruses (**Table 8**). In contrast, IC_50_ values of NBCoV35 and NBCoV63 increased against all mutants by >20-fold and >15-fold, respectively, compared to the activity against the WT virus evaluated in the same cell line. These data suggested that NBCoV compounds may directly interact with K947 and K776.

**Table 8.** Antiviral activity of the NBCoV compounds in a single-cycle assay in A549/AT cells against NL4-3^ΔEnv^-NanoLuc/mutant-SARS-CoV-2 pseudoviruses.

| Compound → | NBCoV35 | NBCoV36 | NBCoV37 | NBCoV63 |
| --- | --- | --- | --- | --- |
| SARS -CoV2 ↓ | IC<sub>50</sub> (nM)<sup>a</sup> |  |  |  |
| WT | 126±3 | 227±18 | 145±12 | 55±3 |
| K947D | 2510±435 | >4000 | >4000 | 1100±25 |
| K947L | 2550±102 | >4000 | >4000 | 896±51 |
| K776D | 3150±312 | >4000 | >4000 | 813±71 |
| K776L | 3587±81 | >4000 | >4000 | 1560±36 |

<sup>a</sup>The reported values represent the means ± standard deviations (n = 3).

### *In vitro* ADME

Assessment of ADME properties in the early stage of drug discovery and development is necessary to reduce drug failure in later stages of drug development^62^. We decided to study the ADME properties of our best inhibitor, NBCoV63, which had potent pan-coronavirus activity, low cytotoxicity, and excellent SI. The ADME properties of NBCoV63 were evaluated by Cyprotex US, LLC (Watertown, MA). Solubility is one of the essential properties of a drug and plays a critical role in drug discovery. The data in **Table 9** indicate that the solubility of NBCoV63 is low. Therefore, special attention should be given to increasing its solubility during its optimization phase. One option will be to attach a solubilizing group in a non-pharmacophoric site which is expected to increase its solubility. The apparent permeabilities of NBCoV63 in the Caco-2 permeability assay are also low. This assay is used to understand oral drugs’ gastrointestinal (GI) absorption. The data also indicate that P-glycoprotein (P-gp)-mediated efflux is involved. The primary role of P-gp is to protect the body from harmful substances by removing drugs absorbed in the intestines back into the gut lumen. We examined the metabolic stability of NBCoV63 in the human liver microsome because the liver is the most important site of drug metabolism in the body. The clearance data (Cl_int_) in **Table 9** indicate that NBCoV63 is a low-clearance compound with a half-life of >180 min. It is worthwhile to mention that low clearance is often a goal in drug development because it allows for lower doses to be used, which reduces drug-related toxicities. The binding data for NBCoV63 in human plasma showed that the inhibitor is >99.5% bound (**Table 9**), which is high. Although many drugs have >98% plasma protein binding, high protein binding does not affect the success of drug candidates. The misconception that high plasma protein binding is a problem for drug candidates was elegantly discussed by Smith et al. in 2010^63^. To identify potential drug–drug interactions, we evaluated NBCoV63 against six CYP450 isoforms (CYP1A2, CYP2B6, CYP2C8, CYP2C9, CYP2C19, CYP2D6) that play an essential role in metabolizing almost 80% of all drugs in the human body^64, 65^.

**Table 9.** *In vitro* ADME profile of one of the most potent inhibitors NBCoV63

| Assay performed | *in vitro* ADMET | Inhibitor |
| --- | --- | --- |
|  |  | NBCoV63 |
| Solubility (µM) | Phosphate buffer, pH7.4 | 6.25 |
| Caco-2 permeability (mean P<sub>app</sub>, x 10<sup>-6</sup> cm/sec) | A-to-B | 0.00242 |
|  | B-to-A | 0.19 |
|  | Efflux Ratio | 42.3 |
| Metabolic stability (human liver microsomes) | Cl<sub>int</sub> (µl/min/mg protein) | 3.4 |
|  | Half-life (min) | >180 |
| Protein binding (human plasma) | % bound | >99.5 |
| Cytochrome P450 inhibition, IC<sub>50</sub> (µM) | CYP1A2 (a-Naphthoflavone) | >25 |
|  | CYP2B6 (Ticlopidine) | >25 |
|  | CYP2C8 (Quercetin) | >25 |
|  | CYP2C9 (Sulphaphenazole) | >25 |
|  | CYP2C19 (Ticlopidine) | >25 |
|  | CYP2D6 (Quinidine) | > 25 |

The following guideline is usually used for the CYP inhibition assessment^66^:

IC50 > 10 μM (CYP inhibition low)

< 10 μM (CYP inhibition moderate)

< 3 μM (CYP inhibition high)

This classification indicated that NBCoV63 had no inhibition up to 25 μM (**Table 9**), indicating high tolerance to CYP-mediated metabolism.

## CONCLUSIONS

We have used the pharmacophoric knowledge from a set of our earlier reported benzoic acid derivatives, which contained ene-rhodanine moiety, known as a promiscuous “frequent hitter” in high-throughput screening, as potent pancoronavirus inhibitors. To our knowledge, this is the first example of using the information of the pharmacophores from PAINS-containing compounds to identify a new set of inhibitors devoid of any PAINS. We identified a series of small molecule benzoic acid analogs that showed highly potent pancoronavirus activity in a pseudovirus-based single-cycle assay. These inhibitors also showed low cytotoxicity, therefore, exhibiting high SI values. One of them, NBCoV63, showed the most consistent high potency against SARS-CoV-2, SARS-CoV, and MERS-CoV, their mutants, and VOCs in different cell lines. Most significantly, NBCoV63 also showed similar efficacy profiles to Remdesivir against an authentic SARS-CoV-2 (Hong Kong strain) and its Delta variant; however, Remdesivir showed about 2-fold better activity against the Omicron variant in a plaque assay in Calu-3 cells. In addition, we showed that NBCoV63 inhibited virus-mediated cell-to-cell fusion in a dose-dependent manner. The in vitro ADME data of NBCoV63 show drug-like properties. However, the solubility of this class of inhibitors needs to be improved. Nevertheless, NBCoV63 can be considered a lead candidate for medicinal chemistry-based optimization.

## EXPERIMENTAL SECTION

### Cells and plasmids

The MRC-5 (fibroblasts cell line isolated from lung tissue), Caco-2 (epithelial cell line isolated from colon tissue), Calu-3 (epithelial cell line isolated from lung tissue), and HEK293T/17 cells were purchased from ATCC (<u>Manassas, VA</u>). The human lung carcinoma A549 cells expressing human ACE2 & TMPRSS2 and the spike pseudotyping vectors Delta Variant (B.1.617.2) pLV-SpikeV8, Omicron Variant (B.1.1.529/BA.1) pLV-SpikeV11) and Omicron Variants (BA.4/BA.5) pLV-SpikeV13) were purchased from InvivoGen (San Diego, CA). The human T-Cell lymphoma Jurkat (E6-1) cells were obtained through the NIH ARP. The 293T/ACE2 cells and the two plasmids pNL4-3^ΔEnv^-NanoLuc and pSARS-CoV-2-S^Δ19^ were kindly provided by Dr. P.Bieniasz of Rockefeller University^67^. The pSV-A-MLV-Env (envelope) expression vector ^68, 69^ and the Env-deleted proviral backbone plasmids pNL4-3.Luc.R-E-DNA ^70, 71^ were obtained through the NIH ARP. The two plasmids, pSARS-CoV, and pMERS-Cov, were kindly provided by Dr. L. Du of the New York Blood Center. The expression vector containing the SARS-CoV-2 full Spike wild-type (WT) gene from Wuhan-Hu-1 isolate (pUNO1-SARS-S) was purchased from InvivoGen (San Diego, CA). The pFB-Luc plasmid vector was purchased from Agilent Technologies (Santa Clara, CA).

### Small molecules

We screened a set of twenty-two small molecules, of which NBCoV35-37 were purchased from Chembridge Corporation (San Diego, CA) and NBCoV62-82 were purchased from ChemDiv, Inc (San Diego, CA). The purity of all purchased compounds is >90%. Details of the analyses are reported in the Supporting Information.

### Pseudoviruses preparation

Pseudoviruses capable of single-cycle infection were prepared by transfecting 8×10^6^ HEK293T/17 cells with a proviral backbone plasmid and an envelope expression vector by using FuGENE HD (Promega, <u>Madison, WI</u>) and following the manufacturer’s instructions as previously described^46^. We used the HIV-1 Env-deleted proviral backbone plasmid pNL4-3^ΔEnv^-NanoLuc DNA and the Envs SARS-CoV-2 and its VOCs, SARS-CoV and the MERS-CoV to obtain the respective pseudoviruses. For the A-MLV pseudovirus, we used the Env-deleted proviral backbone plasmids pNL4-3.Luc.R-.E-DNA and the pSV-A-MLV-Env expression vector. Pseudovirus-containing supernatants were collected two days after transfection, filtered, tittered, and stored in aliquots at −80 °C. Pseudovirus titers were determined with the Spearman-Karber method^72^ to identify the 50% tissue culture infectious dose (TCID_50_) by infecting the different cell types as previously described^46^. The incorporation of the respective spike proteins into the pseudoviruses was confirmed as previously described^46^.

### Measurement of antiviral activity

The antiviral activity of the NBCoV small molecules was evaluated in a single-cycle infection assay by infecting different cell types with the SARS-CoV-2 and its VOCs, SARS-CoV, or MERS-CoV pseudoviruses as previously described ^32, 46, 73^.

### A549/AT cells

We evaluated the antiviral activity by preincubating aliquots of the pseudovirus SARS-CoV-2, its VOCs, and SARS-CoV (1500 TCID_50_/well and at an MOI of 0.1) with the escalating concentrations of the NBCoVs for 30 minutes before adding to the A549/AT cells (1×10^4^ cells /well in a 96-well cell culture plate). A549/AT cells cultured with medium with or without pseudoviruses were included as positive and negative controls, respectively. Following 48 h incubation, the cells were washed with PBS and lysed with 50 μL of lysis buffer (Promega). Twenty-five μL of the lysates were transferred to a white plate and mixed with the same volume of Nano-Glo® Luciferase reagent. The luciferase activity was measured with the Tecan SPARK. The percent inhibition and the IC_50_ (the half-maximal inhibitory concentration) values were calculated using the GraphPad Prism 9.0 software (San Diego, CA).

### 293T/ACE2 cells

The 96-well plates used for the 293T/ACE2 cells were coated with 50 μL of poly-l-lysine (Sigma-Aldrich, St. Louis, MO) at 50 μg/mL as previously described^46^. The neutralization assay and the data collection were performed as described above for A549/AT cells.

### Calu-3 cells

The antiviral activity of the small molecules in Calu-3 cells was evaluated as reported above by plating 5×10^4^ cells/well the cells in a 96-well cell culture plate and incubating overnight. On the following day, aliquots of the pseudovirus SARS-CoV-2 or the VOC Omicron (BA.4/BA.5) at about 7000 TCID_50_/well at an MOI of 0.1 were pre-incubated with escalating concentrations of the NBCoV small molecules for 30 minutes then added to the cells. The assay and data collection were performed as described above.

### Caco-2 and MRC-5 cells

Caco-2 and MRC-5 cells were plated at 1×10^4^ cells/well in a 96-well cell culture plate and incubated overnight. On the following day, aliquots of the MERS-CoV pseudovirus at about 1500 TCID_50_/well at an MOI of 0.1 were pretreated with graded concentrations of the small molecules for 30 min and added to the cells. The assay and data collection were performed as described above.

### SARS-CoV-2 plaque reduction assay

We evaluated NBCoV63 and the control article Remdesivir by using a traditional plaque reduction assay. The assay was performed in a 24-well plate using Calu-3 cells. The cells were seeded 24 h before the assay at a density to reach confluency. The test compound and the control article were diluted 1:2 at a 2x concentration in a 96-well plate in duplicate. The final test compound concentration range was from 0.04 μM to 20 μM, and the final control article concentrations range was from 0.1 μM to 50 μM. An equal volume of the diluted virus was added to each well to make the final compound concentration 1x. The compounds were tested against the following viruses: SARS-CoV-2 Hong Kong/VM20001061/2020 (HK) at 0.5 MOI, SARS-CoV-2 Omicron hCoV-19/USA/NY-MSHSPSP-PV56475/2022, and SARS-CoV-2 Delta hCoV-19/USA/PHC658/2021 at 0.01 MOI. Twenty-four well cell seeding at 240,000 cells/well was used for MOI calculations. All incubations were done in a 37°C and 5% CO_2_ incubator in an approved BSL3 laboratory. The virus and compound dilutions were gently mixed and incubated for 1 h. Following incubation, the virus and diluted compound mixtures were transferred to the respective wells of the Calu-3 cells. The mixture was incubated with the Calu-3 cells for 1 h with gentle rocking every 15 minutes. Following incubation to each well, a 500 μl agarose plug (1:1 2x media in 1% agarose) mixture was added. The agarose plug was allowed to solidify before placing it in the incubator for 72 h. Following incubation, 500 μl of 10% formalin was added to the agarose plug for at least 1 h. The formalin and agarose plug was removed, and the wells were stained with 1% crystal violet in 10% formalin for 15 minutes. The wells were washed twice, and the plaques were counted to determine the IC_50_. Compounds without virus were tested for toxicity using the same procedure, and the CC_50_ was calculated. Prism software was used for all calculations.

### Evaluation of cytotoxicity

The cytotoxicity of NBCoV small molecules in the different cell types was performed in parallel with the neutralization assay and evaluated using the colorimetric CellTiter 96® AQueous One Solution Cell Proliferation Assay (MTS) (Promega, <u>Madison, WI</u>) following the manufacturer’s instructions. Briefly, for the cytotoxicity assay in A549/AT, 293T/ACE2, Caco-2, and MRC-5 cells, 1×10^4^/well cells were plated in a 96-well cell culture plate and incubated overnight; for the same assay in Calu-3 cells, we used 5×10^4^/well cells. The following day, aliquots of escalating concentrations of the NBCoV compounds were added to the cells and incubated at 37 °C. Following 48 h incubation, the MTS reagent was added to the cells and incubated for 4 h at 37 ºC. The absorbance was recorded at 490 nm. The percent of cytotoxicity and the CC_50_ values were calculated using the GraphPad Prism 9.0 software.

### Drug sensitivity of spike-mutated pseudovirus

Amino acid substitutions or deletions were introduced into the pSARS-CoV-2-S^Δ19^ expression vector by site-directed mutagenesis (Stratagene, La Jolla, CA) by following the manufacturer’s instructions and using mutagenic oligonucleotides SaCoV2-K417T-F: GCTCCAGGGCAAACTGGA<u>ACG</u>ATTGCTGATTATAATTAT and SaCoV2-K417T-REV: ATAATTATAATCAGCAATCGTTCCAGTTTGCCCTGGAGC; SaCoV2-K947L-F: ACAGCA AGCGCCCTGGGATTGCTGCAGGACGTGGTCAAC and SaCoV2-K947L-REV: GTTGACCACGTCCTGCAGCAATCCCAGGGCGCTTGCTGT; SaCoV2-K947D-F: ACAGCAAGCGCCCTGGGAGATCTGCAGGACGTGGTCAAC and SaCoV2-K947D-REV: GTTGACCACGTCCTGCAGATCTCCCAGGGCGCTTGCTGT; SaCoV2-K776L-F: ATCGCCGTGGAACAGGACTTGAACACCCAAGAGGTGTTC and SaCoV2-K776L-REV: GAACACCTCTTGGGTGTTCAAGTCCTGTTCCACGGCGAT; SaCoV2-K776D-F: ATCGCCGTGGAACAGGACGACAACACCCAAGAGGTGTTC and SaCoV2-K776D-REV: GAACACCTCTTGGGTGTTGTCGTCCTGTTCCACGGCGAT. Primers for mutations E484K, N501Y, and D614G have already been described ^46^. Site mutations were verified by sequencing the entire spike gene of each construct. To obtain the SARS-CoV-2 pseudovirus carrying the amino acid substitutions, HEK293T/17 cells were transfected with the HIV-1 Env-deleted proviral backbone plasmid pNL4-3^ΔEnv^-NanoLuc DNA and the mutant pSARS-CoV-2-S^Δ19^, as described above. Pseudoviruses were tittered by infecting 293T/ACE2 and A549/AT cells, as described above. A549/AT and 293T/ACE2 cells were infected with the ENV-mutated pseudoviruses, pretreated for 30 min with different concentrations of the NBCoVs and incubated for 2 days to measure the activity of the compounds against the pseudoviruses expressing different point mutations, as described above. The assay and data collection were performed as described above.

### Cell-to-Cell fusion inhibition assay

For the SARS-CoV-2 mediated cell-to-cell fusion assay, we used Jurkat cells which transiently expressed the luciferase gene and stably expressed the SARS-CoV-2 full Spike wild-type (WT) gene from Wuhan-Hu-1 isolate as donor cells and the 293T/ACE2 as acceptor cells as previously described^46^. Briefly, Jurkat cells were transfected with the SARS-CoV-2 WT expression vector by using FuGene HD and following the manufacturer’s instructions. Following 24 h incubation, the cells were washed and selected for the SARS-CoV-2 spike expression using Blasticidin at 10 μg/mL concentration. In parallel, un-transfected control Jurkat cells were exposed to the same concentration of Blasticidin to rule out cell resistance to the antibiotic; the culture of control cells was entirely depleted in about 14 days. The antibiotic was replaced every four days, and the selection lasted for about 20 days. On the day before the assay, the 293T/ACE2 were plated in a 96-well cell culture plate at 8×10^4^/well, while the Jurkat cells were transfected pFB-Luc expression plasmid DNA. Following 20 h incubation, the Jurkat cells at 8×10^4^/well were incubated with escalating concentrations of the NBCoV compound for 1 h, then transferred to the respective wells containing the 293T/ACE2 cells. Additionally, 293T/ACE2 cells cultured with medium with or without the Jurkat cells were included as positive and negative controls, respectively. As an additional control, a set of 293T/ACE2 cells were incubated with Jurkat cells expressing the luciferase gene only (Jurkat-Luc). The plate was spun for 5 minutes at 1500 rpm and then incubated for 4 h at 37ºC. The wells were carefully washed twice with 200 μL of PBS to remove the Jurkat cells that did not fuse with the 293T/ACE2 cells. Finally, the cells were lysed to immediately measure the luciferase activity to calculate the percentage of inhibition of the SARS-CoV-2 mediated cell-to-cell fusion.

### *In vitro* ADME Study

Details of the *in vitro* ADME study and data analyses can be found in the Supplemental.

## Notes

### Competing Interest Statement

The authors have declared no competing interest.

