## Supplemental Information for "Discovery of highly potent small molecule pan-coronavirus fusion inhibitors"

***In vitro* ADME Study.**

**Microsomal Stability (Intrinsic Clearance)**

| Test Article | Test Conc. | Microsome species | Protein Conc. | Incubation | Analytical Method |
|---|---|---|---|---|---|
| NBCoV63 | 1 μM | Human | 0.5 mg/mL | 0, 5, 15, 30 and 45 min (37°C) | LC- MS/MS |

**Experimental Procedure**: The test article was incubated in singlicate with liver microsomes at 37 ºC. The reaction contains microsomal protein in 100 mM potassium phosphate buffer (pH 7.4), 2 mM NADPH, 3 mM MgCl2. A control was run for each test article omitting NADPH to detect NADPH-free degradation. At predetermined time points, as mentioned above, an aliquot was removed from each experimental and control reaction and mixed with an equal volume of ice-cold acetonitrile containing internal standard to stop the reaction and precipitate proteins. The samples were centrifuged to remove precipitated protein, and the supernatants were analyzed by LC-MS/MS to quantify % parent remaining.

**Data Analysis:** Data were calculated as % parent remaining by assuming zero minute time point peak area ratio (analyte/IS) as 100% and dividing remaining time point peak area ratios by zero minute time point peak area ratio. Data were subjected to fit a first-order decay model to calculate slope and, thereby, half-life. Intrinsic clearance was calculated from the half-life and the human liver microsomal protein concentrations using the following equations:

$t_{1/2} = \ln(2) / -k$

$CL_{int} = \ln(2) / (t_{1/2}$ [microsomal protein]) Where

k = slope (elimination constant)

$CL_{int}$ = intrinsic clearance

$t_{1/2}$ = half-life

**Plasma Protein Binding**

| Test Article | Test conc. | Plasma Species | Incubation | Reference Compounds |
|---|---|---|---|---|
| NBCoV63 | 5 µM | Human | 4 hr 37 °C | Warfarin |

**Experimental Procedure**: The test article was added in duplicate to plasma (pH 7.4, ± 0.1, adjusted if necessary). This mixture was dialyzed in a RED device (Rapid Equilibrium Dialysis, Pierce) per the manufacturers' instructions against PBS and incubated on an orbital shaker. At the end of the incubation, aliquots from both plasma and PBS sides were collected and were matrix-matrix matched with an appropriate amount of PBS and blank plasma, respectively. Acetonitrile (three volumes) containing an internal analytical standard (IS) was added to precipitate the proteins and release the test article. After centrifugation, the supernatant was transferred to a new plate and analyzed by LC-MS/MS to obtain peak area ratios (analyte/IS) for determining the fraction unbound.

**Data Analysis:** The extent of binding was reported as a fraction unbound (fu) value which is calculated as detailed below;

fu= PF/PC

Where:

fu = fraction unbound

PC = Test compound in the protein-containing compartment

PF = Test compound in the protein-free compartment.

**Caco-2 Permeability**

| **Test Article** | **Test Conc.** | **Assay Time** | **Transport Direction** | **Reference Compounds** | **Analytical Method** |
|---|---|---|---|---|---|
| NBCoV63 | 10 µM | 2 hours (pH 7.4/7.4) | A→B B→A | Ranitidine Talinolol Warfarin | LC-MS/MS |

**Experimental Procedure:** Caco-2 cells grown in tissue culture flasks was trypsinized, suspended in medium, and the suspensions were applied to wells of a Millipore 96 well plate. The cells were allowed to grow and differentiate for three weeks, feeding at 2- day intervals.

For Apical to Basolateral (A→B) permeability, the test article was added to the apical (A) side and amount of permeation were determined on the basolateral (B) side; for Basolateral to Apical (B→A) permeability, the test article was added to the B side, and the amount of permeation was determined on the A-side. The A-side buffer will contain 100 µM lucifer yellow dye in Transport Buffer (1.98 g/L glucose in 10 mM HEPES, 1x Hank's Balanced Salt Solution) pH 7.4, and the B-side buffer was Transport Buffer at pH 7.4. Caco-2 cells were incubated with test articles in these buffers for 2 hr, and at the end of the assay, donor and receiver side solution samples were collected, quenched by 100% methanol containing an internal standard, and centrifuged at 5000 rpm for 10 minutes at 4ºC. Following centrifugation, the supernatant for donor and receiver side samples was analyzed by LC-MS/MS to determine peak area ratios.

**Data Analysis:**

Data was expressed as permeability (*Papp*):

**Papp = (dQ / dt) / (A x $C_0$)**

Where dQ/dt is the rate of permeation, C0 is the initial concentration of the test article, and A is the monolayer area.

In bi-directional permeability studies, the Efflux Ratio (Re) will also be calculated:

$R_e = P_{app}(B \rightarrow A)/P_{app}(A \rightarrow B)$

Re > 2 indicates a potential substrate for P-gp or other active efflux transporters.

**Turbidimetric Solubility**

| Test Article | Test conc. (μM) | Medium | Incubation | Reference Compounds |
|---|---|---|---|---|
| NBCoV63 | 1.6, 3.1, 6.25, 12.5, 25, 50, 100, 200 μM | PBS (pH 7.4) | 2 hr 37°C | Reserpine<br>Tamoxifen<br>Verapamil |

**Experimental Procedure**: Serial dilutions of the test article were prepared in DMSO at 100x the final concentration. Test article solutions were diluted 100-fold into the buffer in a 96-well plate and mixed. After some time, the presence of precipitate was detected by turbidity (absorbance at

540 nm). An absorbance value > 'mean + 3x standard deviation of the blank' (after subtracting the background) indicated turbidity. For brightly colored compounds, the plate was visually inspected to verify the solubility limit determined by UV absorbance.

**Data Analysis:** The solubility limit was reported as the highest experimental concentration with no evidence of turbidity.

**Microsomal CYP Inhibition Protocol ($IC_{50}$ Determination)**

| Test Article | Test Conc. | CYP Isoforms | Analytical Method |
|---|---|---|---|
| **NBCoV63** | 0.025 - 25 µM | CYP1A2, CYP2B6, CYP2C8, CYP2C9, CYP2C19, CYP2D6, CYP3A4 | LC-MS/MS |

**Experimental Procedure**: The stock solution of the test article was prepared in DMSO at least 400x the top testing concentration and stored at -20 °C. Serial dilutions of the stock solution were prepared in acetonitrile: DMSO for CYP inhibition testing. The final DMSO content in the reaction mixture was equal in all solutions used within an assay and was ≤ 0.25 %. The test article was incubated at seven increasing concentrations in singlicate with pooled human liver microsomes in the presence of 2 mM NADPH in 100 mM potassium phosphate (pH 7.4) containing 5 mM magnesium chloride and a probe substrate. The probe substrate concentration, including tacrine (CYP1A2), bupropion (CYP2B6), amodiaquine (CYP2C8), tolbutamide (CYP2C9), mephenytoin (CYP2C19), dextromethorphan (CYP2D6), midazolam and testosterone (CYP3A4) was optimized as summarized in the table below. The selective CYP inhibitors were screened alongside the test articles as a positive control, including α-naphthoflavone (CYP1A2), ticlopidine (CYP2B6), quercetin (CYP2C8), sulfaphenazole (CYP2C9), ticlopidine (CYP2C19), quinidine (CYP2D6), and ketoconazole (CYP3A4). After optimal incubation at 37 °C (Table 1), the reactions were terminated by the addition of methanol containing internal standard for analytical quantification. The quenched samples were incubated at 4° C for 10 min and centrifuged at 4 °C for 10 min. The supernatant was removed, and the probe substrate metabolite was analyzed by LC-MS/MS.

**Data Analysis:** A decrease in the formation of the metabolite compared to vehicle control was used to calculate an $IC_{50}$ value (the test concentration that produces 50% inhibition).

**Table 1:** Enzyme/Substrate Pairs

| CYP Isoform | Substrate | Substrate Concentration | HLM Concentration | Incubation Time | Positive Control |
|---|---|---|---|---|---|
| CYP1A2 | Tacrine | 5 μM | 0.2 mg/mL | 10 min | α- naphthoflavone |
| CYP2B6 | Bupropion | 100 μM | 0.25 mg/mL | 10 min | Ticlopidine |
| CYP2C8 | Amodiaquine | 5 μM | 0.25 mg/mL | 10 min | Quercetin |
| CYP2C9 | Tolbutamide | 100 μM | 0.5 mg/mL | 15 min | Sulfaphenazole |
| CYP2C19 | Mephenytoin | 100 μM | 0.25 mg/mL | 60 min | Ticlopidine |
| CYP2D6 | Dextromethorphan | 5 μM | 0.5 mg/mL | 10 min | Quinidine |
| CYP3A4 | Midazolam | 2.5 μM | 0.25 mg/mL | 10 min | Ketoconazole |
| CYP3A4 | Testosterone | 50 μM | 0.25 mg/mL | 10 min | Ketoconazole |
